# Phosphocitrate Is Superior to Pyrophosphate in Preventing Soft Connective Tissue Calcification in a Mouse Model of Pseudoxanthoma Elasticum

**DOI:** 10.64898/2026.07.10.737791

**Authors:** Ibtesam Rajpar, Christina Shao, Celina Ng, Fatemeh Niaziorimi, Jacob Beiriger, Petri Turhanen, Koen van de Wetering

## Abstract

Pseudoxanthoma elasticum (PXE) is a rare inherited disorder characterized by progressive ectopic calcification of soft connective tissues, including skin, arteries, and eyes, leading to significant morbidity. PXE results from loss of functional ABCC6, a liver specific ATP efflux conduit. Reduced ATP release into the circulation limits its conversion into AMP and the mineralization inhibitor pyrophosphate (PPi). Consequently, low plasma PPi levels drive ectopic calcification in PXE. Although oral PPi supplementation can inhibit ectopic calcification in Abcc6⁻/⁻ mice, impractically high doses are needed, due to its rapid hydrolysis in the gastrointestinal tract.

Here, we evaluated phosphocitrate, an exceedingly more potent mineralization inhibitor, in vitro and in Abcc6⁻/⁻ mice. In ATDC5 cells, 1 µM phosphocitrate significantly inhibited mineralization following induction, comparable to approximately tenfold higher concentrations of PPi. In vivo, daily intraperitoneal administration of phosphocitrate (4.7 µmol/kg bw) markedly reduced calcification in muzzle skin and kidneys, whereas an at least fivefold higher dose of PPi was needed to achieve a similar effect. Oral administration required substantially higher doses of both compounds (∼2.4 mmol/kg bw), but PC remained more effective than PPi at inhibiting soft tissue calcification in Abcc6⁻/⁻ mice.

Importantly, unlike PPi, oral phosphocitrate did not adversely affect skeletal strength or stiffness, even at supraphysiological doses.

In summary, phosphocitrate is a more potent inhibitor of ectopic calcification than PPi in Abcc6⁻/⁻ mice. While optimization of oral delivery remains necessary, its increased potency supports the potential of alternative administration routes, including subcutaneous delivery, as a practical therapeutic strategy for PXE.

## INTRODUCTION

Pseudoxanthoma elasticum (PXE) is a rare, autosomal recessive disorder characterized by the deposition of calcium phosphate composites primarily in the skin, eyes, and arteries^1^. This connective tissue disorder affects approximately 1 in 50,000 individuals and is of late onset^2^. There currently is no effective or specific treatment available for PXE and the disease therefore slowly progresses after diagnosis. Ectopic calcifications in PXE severely affect quality of life, due to progressive loss of vision, and cardiovascular complications such as intermittent claudication, due to obstruction of the peripheral arteries^3^. Clearly, there is a pressing need for effective treatments that can either block, or effectively remove mineral deposits, to prevent the local and systemic consequences of ectopic calcification prevailing in the soft tissues of PXE patients^4^.

PXE is caused by mutations in the gene encoding the ATP-binding cassette subfamily C member 6 (ABCC6) transporter protein, which is primarily expressed in hepatocytes, where it mediates the efflux of adenosine triphosphate (ATP) into the circulation^5^. In the extracellular environment, ATP is converted to adenosine monophosphate (AMP) and the potent endogenous mineralization inhibitor pyrophosphate (PPi), by the ectonucleotidase ectonucleotide pyrophosphatase/phosphodiesterase 1(ENPP1)^6^. Absence of ABCC6 activity in PXE patients results in reduced levels of PPi in the blood circulation and, consequently, soft tissue calcification^7^. Hence, the increased ratio of inorganic phosphate (Pi) to PPi in soft tissues is often used to explain pathological calcification in PXE^7^.

Since reduced levels of circulating PPi underlie the clinical manifestations in PXE patients, recent efforts by our group as well as others have focused on the development of strategies to increase circulating concentrations of this mineralization inhibitor^7–9^. Because of its short half-life, any PPi therapy for PXE would require daily, lifelong, treatment. This makes the oral route the preferred method for PPi administration to PXE patients. In fact, at the time of writing this paper, a clinical trial for the treatment of PXE patients with a PPi formulation for oral administration was underway in France, of which the results will become available in 2027^10^. In this regard, our group as well as others have incorporated the Abcc6-/- mouse, a validated animal model for PXE. Importantly, Abcc6-/- mice suffer from calcification in the skin, eyes and blood vessels, due to reduced levels of circulating PPi^1^, thereby closely mimicking human PXE. Several translational studies have focused on administration of PPi to Abcc6-/- mice by the oral route, with the goal of increasing plasma levels of this metabolite, and thereby reducing ectopic calcification ^7,9^. We recently found that only very high doses of orally administered PPi, doses impractical for human use, blocked disease progression in Abcc6-/- mice ^8^. Further, the administration of PPi by the alternative subcutaneous route may cause undesirable side effects, such as skin necrosis^11^. The clinical applicability of PPi substitution therapy for PXE is therefore not straightforward, prompting the development of alternative treatment strategies, perhaps involving more potent mineralization inhibitors, allowing for lower drug dosage, frequency of administration and/or to increase drug uptake in the gut.

Given the limitations of oral PPi substitution therapy to treat PXE, we set out to test phosphocitrate (PC), an exceedingly more potent mineralization inhibitor than PPi, for its potential to prevent ectopic calcification in Abcc6-/- mice. PC, originally identified in the seventies in mitochondrial extracts from rat livers^12^, was postulated to protect delicate mitochondrial membranes from the detrimental effects of calcium phosphate crystallization^13^. More recently, Cheung *et al.* have provided evidence to support an inhibitory role of PC in the intraarticular mineralization of aging guinea pigs, a model known for the development of spontaneous osteoarthritis in the knee joints^14^. Further, in 3-4 week old mice with progressive ankylosis (*ank/ank*), PC treatment was associated with increased forelimb joint mobility following 20 days of treatment^15^. PC has been shown to directly bind amorphous calcium phosphate to prevent the formation of hard hydroxyapatite crystals^16,17^. PC has also been shown to potently inhibit calcium phosphate precipitation in cell-free systems^17^. However, the effects of PC on the mammalian skeleton and skeletal functions are also not well understood. Cheung *et al.* noted no differences in the degree of subchondral bone mineralization between PC-treated and saline-treated guinea pigs, suggesting that PC may not adversely affect bone^14^. Nonetheless, a thorough evaluation of the effects of PC on bone tissue is lacking.

To the best of our knowledge nothing is known about the anti-calcifying potential of PC in soft connective tissues or hard mineralized bone, especially in those affected by PXE. In this study, we tested the hypothesis that PC is a more powerful ectopic calcification inhibitor of soft connective tissues in Abcc6-/- mice than PPi. As an initial proof-of-concept experiment, we first determined whether increasing concentrations of each inhibitor affected the mineralization of ATDC5 cells *in vitro*. Next, Abcc6-/- mice received incremental doses of PC or PPi, administered by daily i.p. injection. Soft tissue calcification was quantified by standard µCT densitometry. Finally, to assess PC potency by the oral route, Abcc6-/- mice were treated with incremental doses of PC or PPi by daily oral gavage. To identify the potential adverse effects of these drug treatments on the skeleton, structural and mechanical properties of the femora were assessed. We expect results from these studies to benefit future investigations of treatments and clinical trials for PXE, and other diseases of ectopic calcification.

## MATERIALS AND METHODS

### Synthesis of phosphocitrate

PC was prepared by multistep synthesis procedure described elsewhere^18^. The final product was characterized by necessary nuclear magnetic resonance (NMR) techniques, and the purity was > 96% in all prepared batches according to ^1^H and ^31^P NMR spectra.

### Mineralization assay

To study effects of PC and PPi on mineralization *in vitro* we employed ATDC5 cells, as previously described^5^, with modifications: cells were cultured for 4 days in cell propagation medium comprising of DMEM/F12 (50/50), 5% fetal calf serum (FCS), 100 U/ml pen/strep solution, 10 µg/ml transferrin and 30 nM sodium selenite (all Invitrogen). Next, after confluent monolayers had formed, cells were cultured for an additional 7 days in differentiation medium comprising of DMEM/F12 (50:50) with 1.5% FCS, 100 U/ml pen/strep solution, insulin-transferrin-sodium selenite-ethanolamine solution (ITS, 1:100), and ascorbic acid (50 µg/ml). Next, cells were cultured for 48 hours in mineralization medium consisting of alpha MEM, containing 1.5% FCS, 8 mM phosphate, 100 U/ml pen/strep, 10 µg/ml insulin, 5.5 ug/ml transferrin, 6.7 ng/ml sodium selenite and 2 µg/ml ethanolamide and 24 mM HEPES (pH 7.4), with or without mineralization inhibitor at the indicated concentration. To quantify mineral deposition on the ATDC5 monolayers, cells were fixed in 70% ethanol, washed twice with PBS and stained with Alizarin Red S^5^. Alizarin Red S-bound calcium phosphate precipitates were subsequently solubilized in 10% cetylpyridinium chloride in 10 mM phosphate buffer, and absorbance at the 550nm wavelength was determined using a microplate reader (Tecan Spark).

### Animals

*Abcc6*-/- mice used in this study were on a C57BL/6J background^19^, received food and water ad libitum, and were housed in specialized rodent maintenance facilities with a 12-hour light/12-hour dark cycle. To accelerate the development of ectopic mineral deposits, animals were fed a special ‘acceleration diet’ (TD.00442) containing 0.4% calcium, 0.85% phosphate, 0.04% magnesium and 100 IU/g vitamin D directly following weaning until euthanasia. We first studied the efficacy of PC to prevent ectopic calcification after its daily i.p. administration. Details of the treatment groups are presented in Table 1. Following 4 weeks of treatment when animals were 7 weeks old, they were sacrificed to collect tissues to assess ectopic calcification in the muzzle skin and kidneys. In a second experiment, we determined the efficacy of PC to prevent ectopic calcification after daily oral gavage feeding. Details of the treatment groups are provided in Table 1. Mice were euthanized following 6 weeks of oral gavage treatment. For both studies, each treatment group had equal numbers of male and female mice (N=5 males, 5 females). Animal studies were approved by the Institutional Animal Care and Use Committee of Thomas Jefferson University, in accordance with the National Institutes of Health Guide for Care and Use of Laboratory Animals, under approval number 22-06-514.

**Table 1:** Experimental and control groups.

**Study 1: daily i.p injections for 4 weeks**
| | Genotype | Inhibitor | Dose (mg/kg bw) | Dose (μmol/kg bw) |
| --- | --- | --- | --- | --- |
| Group 1 | Wild type | none | n.a. | n.a. |
| Group 2 | Abcc6-/- | 0.9% NaCl | 0 | 0 |
| Group 3 | Abcc6-/- | Disodium Phosphocitrate | 0.5 | 1.6 |
| Group 4 | Abcc6-/- | Disodium Phosphocitrate | 1.5 | 4.7 |
| Group 5 | Abcc6-/- | Disodium Phosphocitrate | 5 | 16 |
| Group 6 | Abcc6-/- | Disodium Phosphocitrate | 15 | 48 |
| Group 7 | Abcc6-/- | Disodium Phosphocitrate | 50 | 158 |
| Group 8 | Abcc6-/- | Disodium Pyrophosphate | 5 | 23 |
| Group 9 | Abcc6-/- | Disodium Pyrophosphate | 15 | 68 |
| Group 10 | Abcc6-/- | Disodium Pyrophosphate | 50 | 225 |

**Study 2: daily administration by oral gavage for 6 weeks**
|  | Genotype | Inhibitor | Dose (mg/kg bw) | Dose (mmol/kg bw) |
| --- | --- | --- | --- | --- |
| Group 1 | Wild type | none | n.a. | n.a. |
| Group 2 | Abcc6-/- | 0.9% NaCl | 0 | 0 |
| Group 2 | Abcc6-/- | Disodium Phosphocitrate | 40 | 0.127 |
| Group 3 | Abcc6-/- | Disodium Phosphocitrate | 100 | 0.32 |
| Group 4 | Abcc6-/- | Disodium Phosphocitrate | 250 | 0.8 |
| Group 5 | Abcc6-/- | Disodium Phosphocitrate | 750 | 2.4 |
| Group 6 | Abcc6-/- | Disodium Pyrophosphate | 178 | 0.8 |
| Group 7 | Abcc6-/- | Disodium Pyrophosphate | 530 | 2.4 |

### Blood and tissue collection

Blood samples were collected from anesthetized mice immediately prior to euthanasia as previously described^8,20^. Blood was collected by cardiac puncture, into 1 ml syringes containing 25 µl of an 8.6% tripotassium-EDTA solution. Samples were centrifuged at 4000 rpm (∼1500 rcf) for 10 minutes at 4°C. The upper layer of blood plasma was collected into 1.5 ml tubes and centrifuged at 8000 rpm for 10 minutes to generate platelet-poor plasma. Plasma was stored at -80°C until analysis.

Kidneys and muzzle skin were collected following euthanasia to quantify ectopic calcification. Muzzle skin was stored at -80°C. Kidneys were fixed in 10% formalin and later stored in 70% ethanol. Long bones were collected following euthanasia and stored in PBS-soaked gauze at -80°C.

### Analysis of bone structure and mechanics

Femora were scanned using a Bruker SkyScan 1275 micro computed tomography (µCT) analyzer (Bruker, Billerica, MA) fitted with a 1 mm aluminum filter^21^. Scanning parameters of 55 kV and 181 µA were used to scan bones, with a 74 ms exposure time and a high resolution of 13 microns. Bone scans were reconstructed using nRecon (Bruker) software. Structural bone parameters were quantified using CTan (Bruker) software. For analysis of bone strength and stiffness, a standard three-point bending assay was employed using a material testing system (TA Instrument Electroforce 3200 Series III)^22^. Briefly, femora were placed on custom made fixtures with the condyles facing down. A span length of 7.55 mm was recorded with calipers. Bones were then fractured at the mid-section using a monotonic displacement ramp of 0.1 mm/s. Force and displacement data from the three-point bending test, span length data and a reconstructed µCT image of the cross section of the femur mid-diaphysis were input to a custom GNU Octave script (Octave 6.2.0) to derive material properties of ultimate moment, ultimate stress, bending rigidity and Young’s modulus^23^.

### Quantification of ectopic calcification in soft tissues

The left muzzle skin was stored at -80°C until µCT analysis on a SkyScan 1275, using a 1 mm aluminum filter and 74 ms exposure time (settings described earlier for bone)^21^. Muzzle skin and kidneys were scanned at a resolution of 12 µm. Ectopic mineral deposits in kidneys were quantified using lower cut-off values of 55 kV. Representative images of ectopic mineral deposits in various tissues were generated using DataViewer software (Bruker, Billerica, MA) in combination with Fiji for Maximum Intensity Projection (MIP).

### PPi quantification

PPi in plasma was quantified as described in Jansen *et al.* 2014^20^. PPi incorporated into the mineral phase of bone was quantified as previously described^8,24^. In short, the proximal part of the tibia was removed and bone marrow was subsequently removed by centrifugation at 17000 rpm for 1 minute. Next, marrow-free bones were dissolved in 10% formic acid (1 ml per 25 mg bone) overnight in a shaker-incubator set at 60°C. The next day, solubilized bone extracts were centrifuged at 17000 rpm for 10 minutes to remove organic composites. The supernatant was transferred to a clean tube and diluted in ddH_2_O (1:400 dilution). A standard enzymatic assay of ATP sulfurylase was used to convert PPi to ATP for quantification, as per the method described previously^24^: 2.5 µl of sample was incubated with 77.5 µl of a master mix comprising of 80 µM MgCl_2_, 75 mU/ml ATP sulphurylase (New England Biolabs, Ipswich, MA), 1 µM adenosine 5’ phosphosulphate (APS, Santa Cruz Biotechnology) and 50 mM HEPES buffer (pH 7.4), for 30 minutes at 30° C and 10 minutes at 90° C. ATP was quantified using the SL ATP detection reagent (BioThema, Stockholm, Sweden). Bioluminescence was quantified using a Tecan microplate reader. Sample PPi concentrations were extrapolated from a standard curve of known PPi concentrations and expressed as nmol PPi / mg bone.

For plasma PPi quantification, to 2.5 μl of sample, 80 μl of a mixture containing 75mU/ml ATP sulfurylase, 3µM APS, 80µM MgCl_2_ and 40mM HEPES buffer (pH 7.4) was added. The remaining protocol for plasma PPi quantification was similar to as described above for bone.

### Statistical analysis

For quantification of total mineral content in the muzzles and kidneys, PPi in bone and plasma, and bone structural and mechanical properties, bars in graphical data represent the mean, and error bars represent standard error of mean (SEM). Asterisks represent significant differences compared to Abcc6-/- mice treated with saline. For comparisons of group means, one-way analysis of variance was used with post hoc Dunnett’s tests (p ≤ 0.05). For *in vitro* experiments of mineralization, dots or squares represent the average of technical replicates for each concentration of PC or PPi that was tested, and error bars represent SEM. Statistical analysis was carried out in Prism 11 (Graphpad).

## RESULTS

### Phosphocitrate potently inhibits mineralization of ATDC5 cells in vitro

ATDC5 cells originated from a murine teratoma cell line, and are often used as an *in vitro* model for endochondral ossification^25,26^. When grown in differentiation medium, ATDC5 cells form a dense, collagen-rich extracellular matrix which in the presence of extracellular phosphate is prone to mineralize over time. This provides an excellent cellular *in vitro* model to determine the relative potency of mineralization inhibitors. PC concentrations as low as 1 µM already substantially inhibited mineralization of ATDC5 monolayers. A much higher concentration of 30 µM PPi was needed to inhibit mineralization (Fig 1A). Quantification of mineralization by determining the relative amount of Alizarin Red S attached to the mineral by quantifying the A550, indicated PC had an at least 10-fold higher inhibitory potency than PPi (Figure 1B): 30 µM PPi was needed to completely prevent mineralization, compared to 3 µM for PC. Intriguingly, at concentrations higher than 30 µM, PPi treated ATDC5 cells still evidenced mineralization (Fig 1A & B). We will come back to this unexpected result in the discussion section.

**Fig 1.**
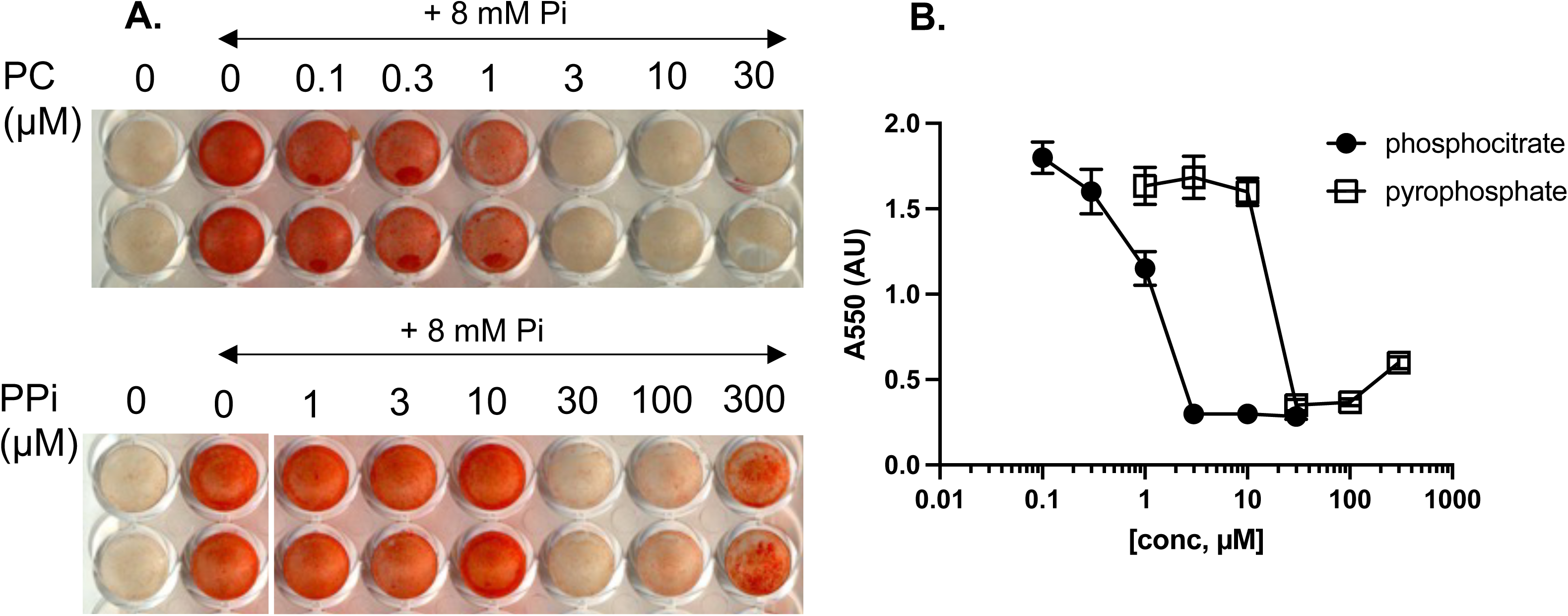
PC inhibits mineralization of ATDC5 cells more potently than PPi in vitro. **A**) Alizarin Red S staining of cells cultured with mineralization media containing PC show that inhibition of mineralization is PC-dose dependent, with complete inhibition at 3 µM. A 10-fold greater dose of PPi was needed to completely inhibit mineralization. **B**) Graphical representation of stain absorbance at A550. Dots/blocks represent mean absorbance for every group, and error bars represent SEM. Group means were compared using one way ANOVA with post hoc Dunnett’s tests (p < 0.05).

### Phosphocitrate is a more potent inhibitor of soft tissue calcification than pyrophosphate in Abcc6-deficient mice

Abcc6-/- mice in this study were fed a diet low in magnesium and high in phosphate to accelerate ectopic calcification^27^. This dietary intervention shortened the treatment period required for the animals and consequently reduced the total amount of PC needed. Under these conditions, 7-week-old Abcc6-/- mice developed robust but variable calcification in both the blood-filled capsules surrounding the vibrissae (muzzle skin calcification) and in the kidneys, consistent with previous reports^8,28^ (Fig. 2). Abcc6-/- mice receiving control treatment (0.9% NaCl) had a mean deposition of mineral in their muzzle skin of 4 x 10^7^ (SD: 4.7 x 10^7^), indicating substantial variability in the extent of mineral deposition (coefficient of variation: 117%). In contrast, wild type mice maintained on the same diet showed no detectable muzzle skin mineralization. Daily intraperitoneal administration of PC for 4 weeks resulted in a dose dependent reduction of muzzle skin mineralization. A low dose of 4.7 μmol/kg body weight (1.5 mg/kg bw) reduced ectopic calcification by approximately 80%, whereas doses > 48 μmol/kg (15 mg PC/kg bw) completely prevented calcification (Fig. 2). In contrast, substantially higher doses of PPi were required to achieve a partial reduction in calcification, and no dose fully prevented mineral deposition. These findings were consistent with our in vitro data, demonstrating the greater potency of PC compared to PPi in inhibiting soft tissue calcification.

**Fig. 2.**
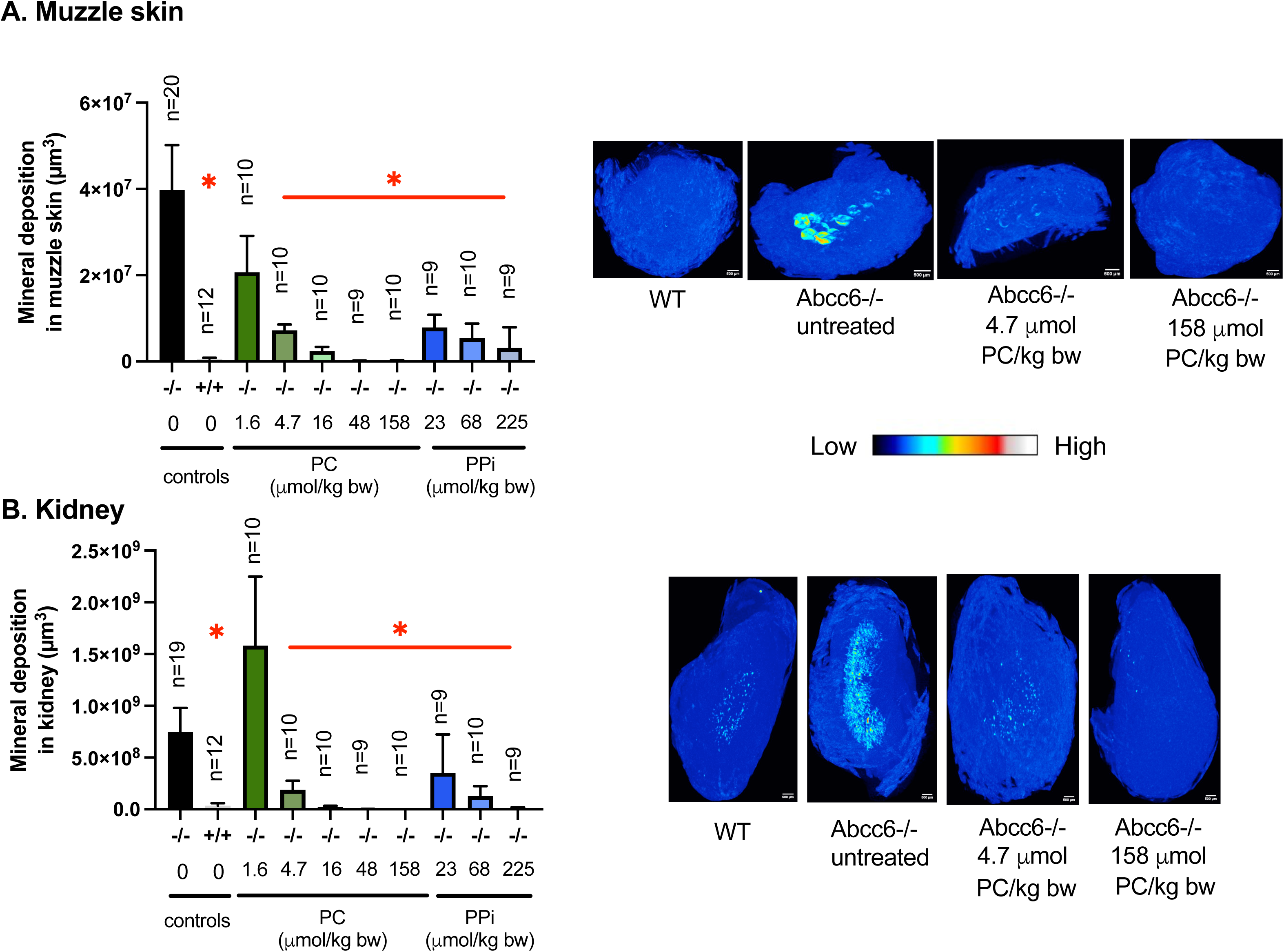
PC is a more potent inhibitor of soft tissue calcification in Abcc6-/- mice than PPi when administered by i.p. injection. Gross examination of microCT sections (images on the right) show presence of mineral deposits (high intensity regions) in the muzzle skin **A**) and kidneys **B**) of untreated Abcc6-/- animals. Significant reduction in mineralization was observed at PC dose of 4.7 µmol/kg bw, and complete inhibition was observed at 48 µmol/kg bw. Images are representative maximum intensity plots of the respective tissue with a mapping of mineral volume fraction. Less inhibition of mineralization was observed at far greater doses of PPi. Group means were compared using one way ANOVA with post hoc Dunnett’s tests. Red asterisks indicate significant differences compared to untreated Abcc6-/- mice, and significance was set at p < 0.05. Error bars represent the SEM. The number of animals in each treatment group is indicated above the corresponding bars, with approximately equal numbers of male and female animals included in each group. Scale bars equal 500 microns.

The effects observed in the muzzle skin were recapitulated in the kidneys following intraperitoneal treatment. Kidney calcifications, primarily localized in the medulla, were markedly reduced in PC-treated Abcc6-/- mice. Even at a low dose of 4.7 μmol/kg bw, PC substantially decreased renal calcification, while higher doses completely abolished it (Fig. 2). As in muzzle skin, higher doses of PPi were required to reduce kidney calcification.

Next, we treated Abcc6-/- mice with PC or PPi by daily oral gavage feeding. Compared to i.p. administration, oral gavage feeding more closely mimics daily oral administration to human PXE patients. It also allowed precise control over treatment parameters including dosing time, dosing intervals, and uniform drug intake across animals. In contrast, administration via drinking water does not allow such control and may result in variable intake, which could explain the limited efficacy of lower doses of PPi reported previously^8^. Abcc6-/- mice received increasing doses of PC (0.13 to 2.4 mmol/kg) or PPi (0.8 and 2.4 mmol/kg) by oral gavage, daily for 6 weeks. Oral PPi administration led to supraphysiological plasma PPi concentrations (Fig. 3C) 10 minutes after dosing. Plasma PPi concentrations reached 11 µM in the 0.8 mmol/kg dosing group and 33 µM in the 2.4 mmol/kg dosing group, compared to 0.23 µM in Abcc6-/- animals receiving saline (Fig. 3C). Ectopic calcification was quantified by µCT densitometry (Fig. 3A, B). Relatively high doses of both PC and PPi (2.4 mmol/kg) were required to fully inhibit muzzle skin calcification when given orally (Fig. 3A). Unexpectedly, PPi at 0.8 mmol/kg had no effect on muzzle skin calcification, despite resulting in supraphysiological PPi plasma concentrations shortly after administration (Fig. 3A, C).

**Fig 3.**
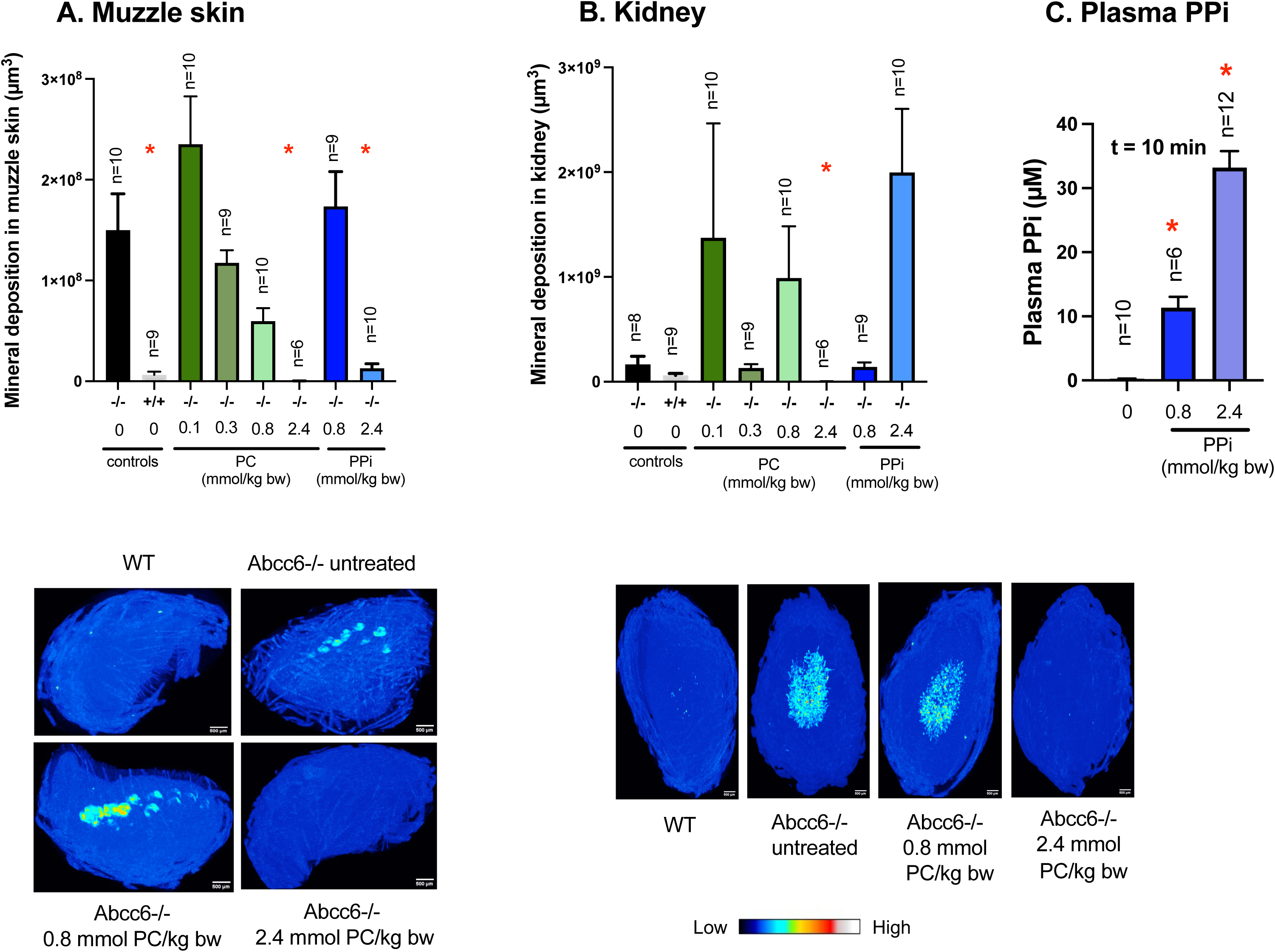
At high doses PC inhibits ectopic mineralization in muzzle skin and kidneys of Abcc6-/- mice after daily oral gavage feeding. Gross examination of microCT sections (images on the right) show presence of mineral deposits (high intensity regions) in the muzzle skin **A**) and kidneys **B**) of untreated Abcc6-/- animals. Almost complete inhibition of mineralization was observed in the muzzle and kidney at PC dose of 2.4 mmol/kg bw. At this dose, supraphysiological concentrations of PPi in the plasma of Abcc6-/- mice were also noted, as quantified using a standard luciferase assay for ATP **C**). Images are representative maximum intensity plots of microCT scans of the respective tissue with a mapping of mineral volume fraction. Group means were compared using one way ANOVA with post hoc Dunnett’s tests. Red asterisks indicate significant differences compared to untreated Abcc6-/- mice, and significance was set at p < 0.05. Error bars represent SEM. Error bars represent the SEM. The number of animals in each treatment group is indicated above the corresponding bars, with approximately equal numbers of male and female animals included in each group. Scale bars equal 500 microns.

For reasons that remain unclear, mineral deposition in kidneys of Abcc6-/- mice maintained on the acceleration diet was highly variable (Fig. 3), making it difficult to detect treatment effects. Control-treated Abcc6-/- mice exhibited pronounced renal calcification, whereas wild-type mice showed substantially lower, though not completely absent, mineral deposition. Despite the substantial variability it was clear that in the kidneys, oral PC - but not PPi - effectively reduced calcification, albeit only at the highest dose tested (2.4 mmol/kg). At this dose, total renal mineral content not only was lower than in saline-treated Abcc6-/- mice, but also than in untreated wild type mice (Fig. 3B).

### Phosphocitrate inhibits ectopic calcification in soft tissues but does not affect skeletal structure and strength in Abcc6-/- mice

To assess the potential adverse effects of oral treatments on the development of mineralized tissues like bone and teeth, we evaluated femoral structure by µCT and bone biomechanical properties by three-point bending assay. Oral PPi treatment was associated with a modest reduction in long bone stiffness, as reflected by decreased bending rigidity and Young’s modulus, the latter accounting for bone geometry (Fig. 4). In contrast, oral administration of PC did not result in similar reductions in bone stiffness. Notably, Abcc6-/- mice treated with 2.4 mmol PC/kg body weight, a dose that effectively inhibited soft tissue calcification, showed no detectable impairment in bone mechanical properties (Fig. 4). Furthermore, no significant differences in cortical or trabecular bone structure were observed between mice treated with either PC or PPi and untreated Abcc6-/- controls (Supp. Fig. 1,2). Finally, the levels of PPi in the mineral phase of tibiae did not differ significantly among treatment groups (Supp. Fig. 3).

**Fig 4.**
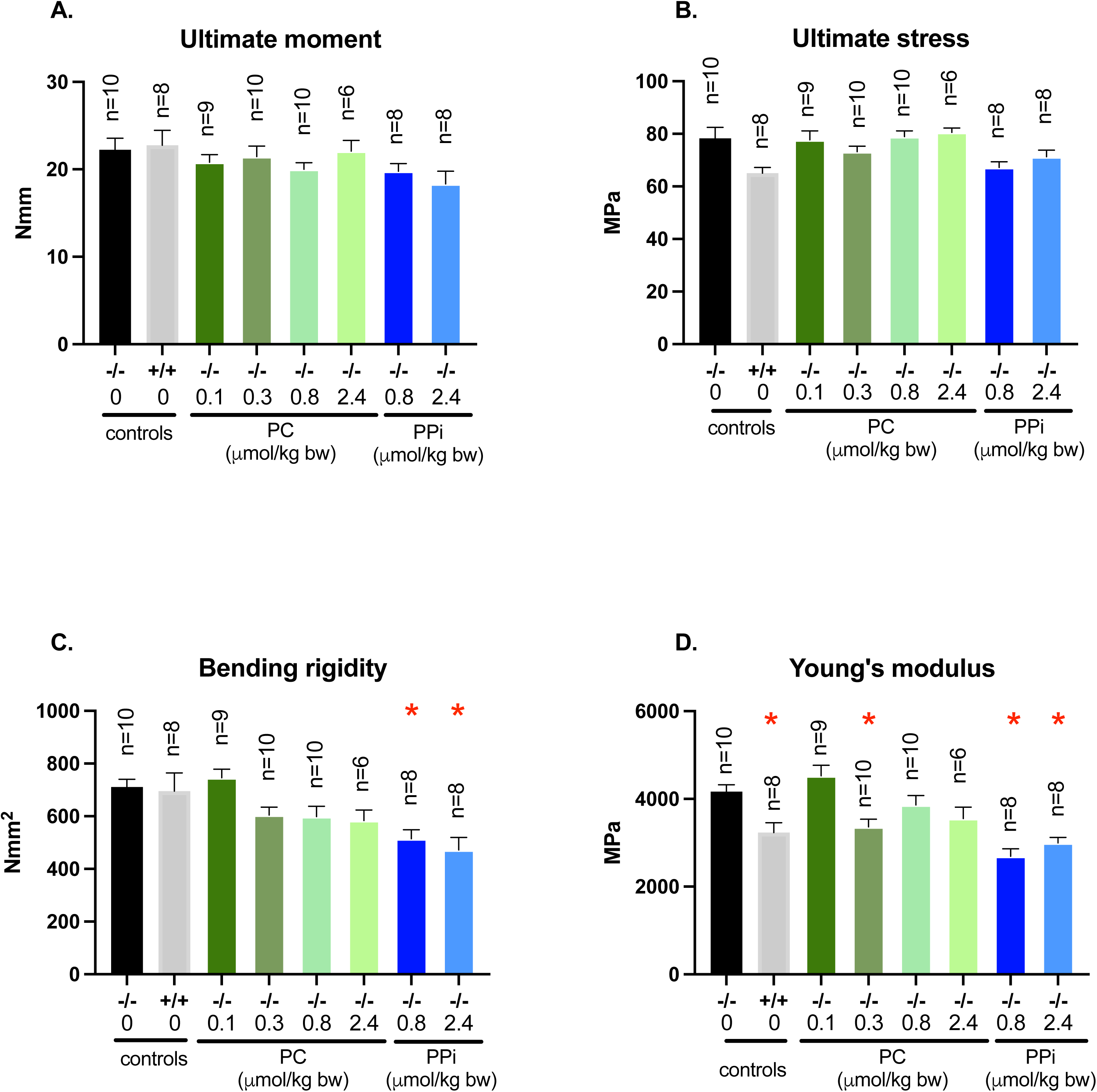
High-dose PC administered via daily oral gavage does not alter the mechanical properties of femora in *Abcc6*⁻/⁻ mice. Various parameters of bone strength and stiffness were quantified using microCT and the three point bending assay. In contrast to PPi, PC did not affect ultimate moment and stress, bending rigidity or Young’s modulus especially at the high 2.4 mmol/kg bw dose which is associated with inhibition of soft tissue calcification. Group means were compared using one way ANOVA with post hoc Dunnett’s tests. Red asterisks indicate significant differences compared to untreated Abcc6-/- mice, and significance was set at p < 0.05. The number of animals in each treatment group is indicated above the corresponding bars, with approximately equal numbers of male and female animals included in each group. Error bars represent SEM.

## DISCUSSION

Phosphocitrate differs structurally from citrate by the presence of an additional phosphate group attached to its tricarboxylic acid backbone^18^. It has been postulated that, by virtue of its greater negative charge, PC can potently bind calcium salts and effectively inhibit the growth and aggregation of calcium-containing salts, such as basic calcium phosphate and calcium pyrophosphate dihydrate salts^29,30^, thereby inhibiting ectopic calcification in soft connective tissues. In a previous study, we demonstrated that only very high, impractical doses of orally administered PPi effectively inhibited ectopic calcification in Abcc6-/- mice^8^. Given that PC is considered a substantially more potent endogenous inhibitor of calcification^31^, we hypothesized that PC would be more effective than PPi *in vivo*. To test this, we compared the efficacy of PC and PPi in preventing ectopic calcification in the Abcc6-/- mouse, a well-established model for PXE. To our knowledge, this is the first direct *in vivo* comparison of these two inhibitors. Our findings demonstrate a markedly higher potency of PC compared to PPi. First, *in vitro* studies using ATDC5 cells showed that PC was approximately tenfold more effective than PPi, with substantially lower concentrations sufficient to completely inhibit extracellular matrix mineralization. Second, *in vivo*, none of the tested i.p. doses of PPi fully prevented calcification of the muzzle skin in Abcc6-/- mice, whereas PC completely inhibited mineral deposition at a dose of 48 µmol/kg. Notably, even a lower dose of 4.7 µmol/kg already resulted in an approximately 80% reduction in calcification. Third, following daily oral administration, high doses of PC completely inhibited calcification in both the muzzle skin and kidneys, whereas comparable doses of PPi were ineffective. Taken together, these results provide strong evidence that PC is a significantly more potent inhibitor of ectopic calcification inhibitor than PPi *in vivo*, regardless of the route of administration. Importantly, oral administration of PC did not adversely affect skeletal properties, even at high doses. Nevertheless, further studies are warranted to optimize the delivery and to enhance the translational potential of PC as a therapeutic agent for PXE.

Our *in vitro* experiments suggest a very high dose of PPi was required to inhibit mineralization, corresponding in efficacy to 1/10^th^ of PC. Both PC and PPi are hydrolyzed primarily by alkaline phosphatases ^32^, which are expressed by ATDC5 cells^26^. However the half-life of PPi in plasma is about 30 minutes, potentially reducing the efficacy of PPi as an anti-mineralization agent^33^. This may explain why PPi does not inhibit matrix mineralization at a low dose of 3 µM. Not much is known of the half-life of PC, although it is inactivated by hydrolysis to inorganic phosphate (Pi) and citrate^34^. At high concentrations, citrate can also inhibit ectopic calcification in Abcc6-/- mice when provided via drinking water^35^. Hence it is possible that differences in degradation of PC and PPi may have contributed to differences in efficacy. Of note, we have tested the inhibitory potential of PC and PPi in the presence of the TNAP inhibitor levamisole. Still, PC was more effective in inhibiting mineralization of ATDC5 cells. These data suggest that increased degradation is not the main reason why PPi is a less effective calcification inhibitor than PC. Interestingly, at very high PPi concentrations of >30 µM, more mineral deposition was seen. Possibly, these high PPi concentrations bind to and stabilize nasent hydroxyapatite crystals.

When administered via drinking water, high doses of PPi are needed to prevent ectopic calcification in Abcc6-/-, as we have shown previously^8^. Here, we attempted to increase the bioavailability of PPi by administering treatment by oral gavage. Compared to drinking water, oral gavage feeding is a controlled method to administer timely doses of PC, and hence may increase drug bioavailability. However, very high impractical doses of PPi and or PC were still needed to block ectopic calcification in Abcc6-/- mice (Fig. 3). In a previous study, a very high dose of 1440 µmol/kg/day (450 mg/kg/day) of PC administered by the oral route in rats, resulted in only a modest 34% inhibition of subcutaneous calcergic plaque formation, following a KMnO_4_ injection in the interscapular region^16^. The efficacy of this dose corresponded to a low i.p. dose of 32 µmol/kg/day^16^. Hence, oral methods of administration may not be practical options for the administration of PC. While no standard methods exist for quantifying PC uptake, these results suggest that negatively charged PC molecules and their hydrophilic nature greatly limit uptake of PC from the gut after oral administration, thereby reducing its inhibitory effects when administered by the oral route. The reduced efficacy of oral PC can be further explained by the presence of abundant phosphatases in the gut that potentially degrade PC^12^. Future studies may investigate optimized parenteral slow-release PC formulations, by using a calcium salt, to circumvent the issues surrounding uptake of PC from the gut.

Based on our data, the lowest effective dose of PC for a healthy adult male when administered by i.p. injection would be ∼0.1 grams PC per day. However, i.p. injection is not feasible in human patients. The pharmacokinetics of drugs administered by subcutaneous injection are very similar to i.p. injection. The subcutaneous route may therefore be convenient for daily PC administration. Notably, other drugs are commonly administered subcutaneously, such as the anticoagulant enoxaparin, which is given by subcutaneous injection at a dose of 1.5 mg/kg. A similar dose of PC is required to inhibit 80% calcification in muzzles of Abcc6-/- mice (4.7 µmol/kg).

Substantial variability in total mineral content was observed in both kidneys and muzzle skin within the same treatment groups. Six-week treated Abcc6-/- control animals in the oral gavage study exhibited more than a threefold increase in muzzle skin mineralization compared to animals treated by i.p. injection for 4 weeks. The initial conclusion we drew from this result is that if oral gavage treatment lasted for 4 instead of 6 weeks, we may have not seen this increase. However, the opposite trend was observed in the kidneys: control-treated Abcc6-/- mice showed over threefold higher renal mineral deposition following i.p. treatment compared to oral gavage feeding. Within the treatment groups, route-dependent differences were also apparent. When administered orally, PC appeared to increase renal mineral content at the lowest tested dose (0.13 mmol/kg) dose. In contrast, the similar dose of PC (158 µmol/kg) when delivered by i.p. injection, completely prevented kidney calcification. As noted earlier, oral PC administration may be less effective in treating kidney calcification, due to hydrolysis of PC to citrate and Pi, and increased accumulation of Pi which may stimulate ectopic calcification. However, these trends did not reach statistical significance and should be interpreted cautiously, as they are strongly influenced by the high inter-animal variability observed in ectopic calcification in Abcc6-/- mice on acceleration diet. The source of this variability remains unclear, but may reflect differences in the microenvironment of mice, social hierarchy or epigenetic factors among mice. For example, in mice with progressive ankylosis, while the underlying cause of joint calcification is genetic, the actual disease penetrance may depend heavily on the mechanical loading of the joints, due to differences in cage running activity^37^. Notably at the highest dose tested (2.4 mmol/kg body weight), consistent daily oral administration of PC completely inhibited kidney calcification, whereas orally administered PPi failed to reduce renal calcification in Abcc6-/- mice on acceleration diet^8^.

In the present study we used the aforementioned acceleration diet, which is a low magnesium, high phosphate formulation that promotes soft tissue calcification in Abcc6-/- mice^27^. We acknowledge that this diet may not fully recapitulate the late onset and slowly progressive nature of ectopic calcification in PXE patients. However, its use was necessitated by the limited availability of PC, which required us to shorten the duration of treatment. Importantly, use of the acceleration diet also resulted in increased variability in soft tissue calcification compared to Abcc6-/- mice maintained on standard chow. This elevated variability reduces statistical power, necessitating use of larger animal cohorts to detect treatment effects. In addition, Abcc6-/- mice on the acceleration diet developed kidney calcifications, a phenotype not observed in animals maintained on standard chow. An alternative to using the acceleration diet, is the Abcc6-/-;*Ank^ank/wt^* double mutant mouse model of ectopic calcification, recently developed in our laboratory (unpublished). These mice lack *Abcc6* expression and have only one functional copy of the *ank* gene. Compared to single mutant Abcc6-/- mice, the double mutants exhibit markedly more severe muzzle skin calcification, yet do not display the pronounced ankylosis phenotype seen in animals homozygous for the inactivating *ank* mutation. This model may therefore provide a useful platform for future studies without the need for a dietary acceleration strategy.

A limitation of this study is the lack of a straightforward method to quantify PC levels in the circulation following treatment. At present, uptake and bioavailability can only be assessed indirectly through measurements of mineral deposition. Finally, due to limited access to PC, we were unable to optimize its formulation. For example, alternative form such as calcium salts of PC^14^, which may improve intestinal absorption and enhance efficacy, were not evaluated in this study. Another option to boost the uptake of hydrophilic drugs after oral administration is by their physical encapsulation within lipid nanoparticles^38^.

Several experimental treatments for PXE are currently in development like ENPP1 enzyme substitution therapy, bisphosphonates, and TNAP inhibitor therapy^6,39–41^. However, these treatments may have limited inhibitory effects on ectopic calcification, can be expensive, and may produce undesirable side effects. Here, we show that PC potently inhibits ectopic calcification when administered by the i.p. route, however, impractical doses of drug were needed for oral administration. Further, one high dose of PPi/day may not effectively prevent calcification, and results of the current trial on oral PPi may better inform translational studies in the future^10^. Future studies will also aim to further develop PC formulations and methods of administration. For example, subcutaneous injection may be an attractive treatment route to increase drug bioavailability, however, whether daily administration of an effective dose of PC is feasible and safe, is yet to be investigated. Further, strategies to combine the inhibitors PC and PPi or PC and magnesium for synergistic effects on calcification inhibition in Abcc6-/- mice, will help identify the most promising therapies for PXE.

## ACKNOWLEDGEMENTS

This research was supported by the National Institutes of Arthritis and Musculoskeletal and Skin Diseases under award nos. R01AR082460 and R21AR083597 to KW. Further funding for this work was provided by PXE International to KW. The content is solely the responsibility of the authors and does not represent the views of the funding agency.

## AUTHOR CONTRIBUTIONS

Ibtesam Rajpar (Investigation, Methodology, Supervision, Formal Analysis, Writing-original draft, Writing-review and editing), Christina Shao (Investigation, Formal Analysis), Celina Ng (Investigation), and Fatemeh Niaziorimi (Investigation, Formal analysis), Jacob Beiriger (Investigation), and Petri Turhanen (Methodology, Writing-review and editing), KvdW (Conceptualization, Funding acquisition, Project Administration, Methodology, Supervision, Formal Analysis, Writing-original draft, Writing-review and editing).

## DECLARATIONS

The authors declare that they have no known competing financial interests or personal relationships that could have appeared to influence the work reported in this paper.

