## Supplemental figures for "Phosphocitrate Is Superior to Pyrophosphate in Preventing Soft Connective Tissue Calcification in a Mouse Model of Pseudoxanthoma Elasticum"

Supp. Fig. 1: Cortical bone properties of femora in Abcc6-/- mice following oral gavage treatment

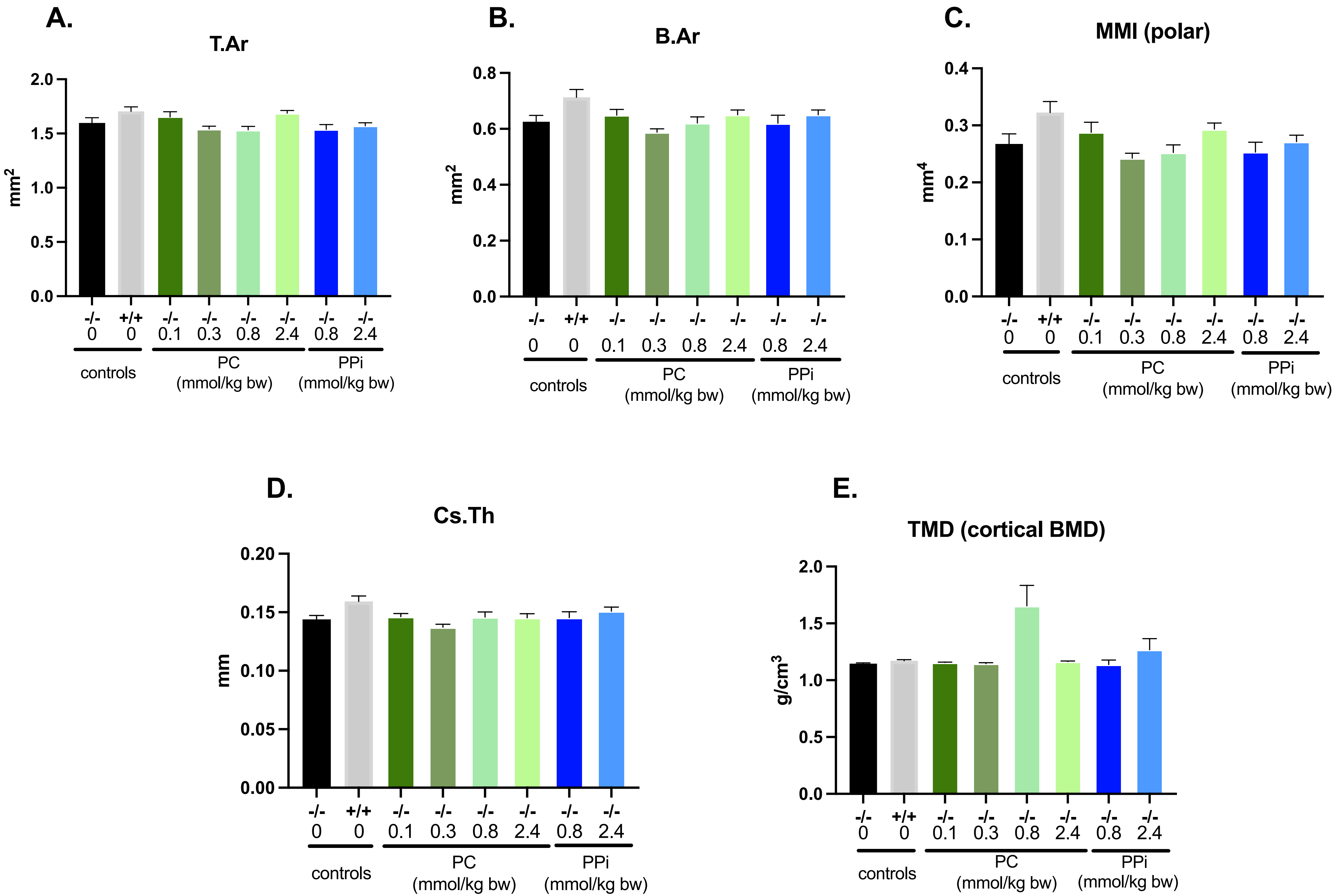

Supp. Fig. 2: Trabecular bone properties of femora in Abcc6-/- mice following oral gavage treatment

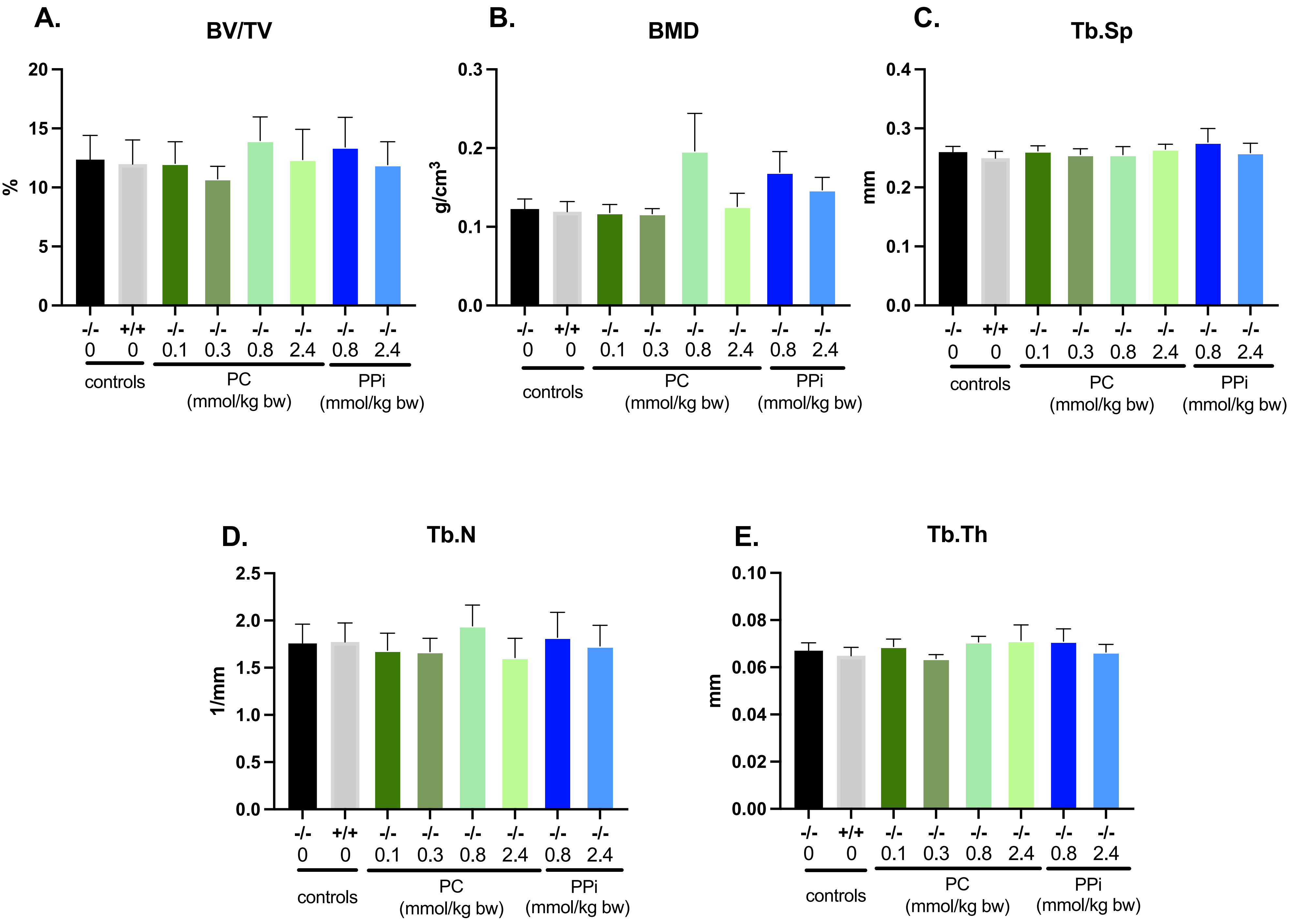

**Supp. Fig. 3: total pyrophosphate in tibiae of Abcc6-/- mice following oral gavage**

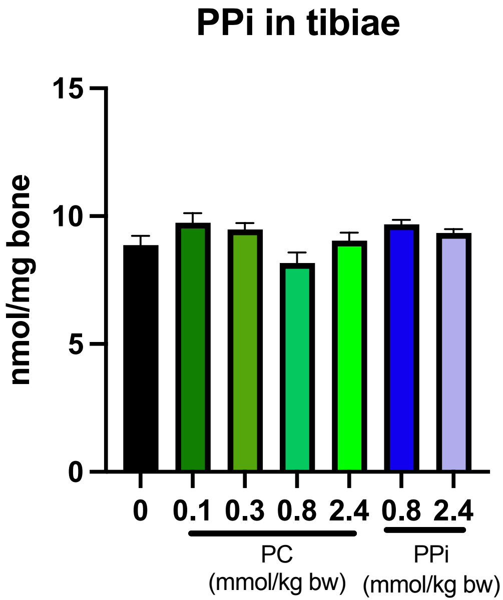

**Supp Fig 1. Effect of oral PC treatment on the cortical bone properties of femora from Abcc6-/- mice.** Various structural parameters of cortical bone were quantified with microCT. TMD (tissue mineral density), Cs.Th (cross sectional thickness), B.Ar. (Bone area), T.Ar. (Tissue area), MMI (mean polar moment of inertia). Group means were compared using one way ANOVA with post hoc Dunnett's tests. Significance was set at $p < 0.05$. Error bars represent the SEM. The number of animals in each treatment group is indicated above the corresponding bars, with approximately equal numbers of male and female animals included in each group. Error bars represent SEM.

**Supp Fig 2. Effect of oral PC treatment on the trabecular bone properties of femora from Abcc6-/- mice.** Various structural parameters of trabecular bone were quantified with microCT. BV/TV (bone volume), BMD (bone mineral density), Tb.Sp (trabecular spacing), Tb.N (trabecular number), Tb.Th (trabecular thickness). Group means were compared using one way ANOVA with post hoc Dunnett's tests. Significance was set at $p < 0.05$. The number of animals in each treatment group is indicated above the corresponding bars, with approximately equal numbers of male and female animals included in each group. Error bars represent SEM.

**Supp Fig 3. Effect of oral PC and PPi treatment on total bone pyrophosphate.** PPi was quantified in tibiae of mice using a standard luciferase assay for ATP. Group means were compared using one way ANOVA with post hoc Dunnett's tests. Significance was set at $p < 0.05$. The number of animals in each treatment group is indicated above the corresponding bars, with approximately equal numbers of male and female animals included in each group. Error bars represent SEM.
